# The circulating lipidome is largely defined by sex descriptors in the GOLDN, GeneBank and the ADNI studies

**DOI:** 10.1101/731448

**Authors:** Dinesh Kumar Barupal, Ying Zhang, Sili Fan, Stanley L. Hazen, W. H. Wilson Tang, Tomas Cajka, Marguerite R. Irvin, Donna K. Arnett, Tobias Kind, Rima Kaddurah-Daouk, Oliver Fiehn

## Abstract

Biological sex is one of the major anthropometric factors which influences physiology, metabolism and health status. We have investigated the effect of sexual dimorphism on the blood lipidome profile in three large population level studies - the Alzheimer’s disease neuroimaging initiative - ADNI (n =806), the GeneBank Functional Cardio-Metabolomics cohort (n= 1015) and the Genetics of Lipid lowering Drugs and Diet Network - GOLDN (n=422). In total, 355 unique lipids from 15 lipid classes were detected across all three studies using LC-MS. Sixty percent of these lipids differed between men and women in all three cohorts, and up to 87% of all lipids demonstrated sex differences in at least one cohort. ChemRICH enrichment statistics on lipid classes showed that phosphatidylcholines, phosphatidylethanolamines, phosphatidylinositols, ceramides, sphingomyelins and cholesterol esters were found at higher levels in female subjects while triacylglycerols and lysophosphatidylcholines were found at higher levels in male participants across the three cohorts. This strong sex effect on the blood lipidome suggests that specific regulatory mechanisms may exist that regulate lipid metabolism in a different manner between men and women. Cohort studies involving blood lipidomics should consider separate analyses for male and female participants instead of combined analyses treating sex as a confounding factor.

## 1. Introduction

Differences between the sexes are one of the fundamental variations in biology[1]. Sex disparities in research can ignore discoveries of the biological mechanisms that are specific to particular sex, leading to missing opportunities in developing new sex-specific therapeutic strategies[2]. Sex differences have been observed for gut microbiota[3] and transcriptome[4], suggesting that sex-specific strategies for health improvement are needed.

Metabolomics has been validated for molecular epidemiology to discover risk factors and biological mechanisms for diseases. Several epidemiological studies have investigated biological sex as a main factor to define a person’s metabolome. Differential network analysis of metabolomics data for 844 healthy subjects suggested a sex-related variability in branched chain amino acids, ketone bodies, and propanoate metabolism [5]. Mittelstrass *et al*. argued that metabolomics analysis in epidemiology should be stratified by sex and showed a strong sex effect in 3,300 participants in the Cooperative Health Research in the Region of Augsburg (KORA) cohort. The study shows that up to 78% metabolites were under sex effect, including amino acids, sphingomyelins, phosphatidylcholines and acyl-carnitines [6]. Metabolomics analysis of 1,756 participants from the KORA F4 study showed that almost 33% of the 507 metabolites were significant different between men and women. The study suggested changes in steroid metabolism, fatty acid, amino acids, purine and dipeptide metabolism differed between the sexes and suggested that a sex-regulated metabolic modules can be identified in the partial correlation network among metabolites [7]. The cross-sectional KarMeN (Karlsruhe Metabolomics and Nutrition) study included 301 participants, yielding a metabolomics dataset that predicted sex descriptors from blood specimen [8]. In a study on 60 subjects it was found that levels of sphingomyelins were higher in women in comparison to men in serum and plasma samples [9]. The study also reported that levels of triacylglycerols were higher in elderly than in younger women. Higher levels of LDL-C, HDL-C, total cholesterol, sphingomyelins and C22:6 fatty acyl-containing phospholipids were observed in women [10]. Similarly, women had higher levels of sphingomyelins and phosphatidylcholines in a French study of 800 participants. In this study, branched chain amino acids and lysophosphatidylcholines were also found to be higher in males [11]. These previous studies highlight the importance of sexual dimorphism in metabolic regulation for lipids and other metabolite levels.

We here report on the effect of sex descriptors on a comprehensive panel of 355 blood lipids from 15 lipid classes in three large cohorts with the largest comparison to date with 2,243 subjects in total. These cohorts included the Genetics of Lipid Lowering Drugs and Diet Network (GOLDN) (*n* = 422), GeneBank, Cleveland Clinic (*n*= 1,015), and the Alzheimer’s Disease Neuroimaging Initiative (ADNI) (*n* = 806). We have used univariate statistics and chemical similarity enrichment analysis to highlight the strong sex effect on the detected lipids.

## 2. Results

### 2.1 Cohort summaries and lipidomics datasets

Table 1 summarizes the cohorts. Participants in the GOLDN cohort were younger compared to the ADNI and GeneBank cohorts. On average, the cohorts consisted of 60% men and 40% women.

**Table 1.** Cohort summaries

| Cohort | Participants | Male | Female | Age |
| --- | --- | --- | --- | --- |
| Alzheimer's Disease<br>Neuroimaging Initiative (ADNI) | 806 | 481 (60%) | 358 | 75.2 ($\pm 6.3$) |
| GeneBank cohort | 1015 | 634 (62%) | 381 | 64 ($\pm 10$) |
| Genetics of Lipid Lowering<br>Drugs and Diet Network | 422 | 207 (49%) | 215 | 49 ($\pm 16$) |

Figure 1 shows the lipidomics data acquisition and processing workflow. All lipidomics data were acquired using identical LC-MS instruments at the West Coast Metabolomics Center, UC Davis. A set of 15 internal standards were added to each sample which were used for retention time correction.

**Figure 1.**
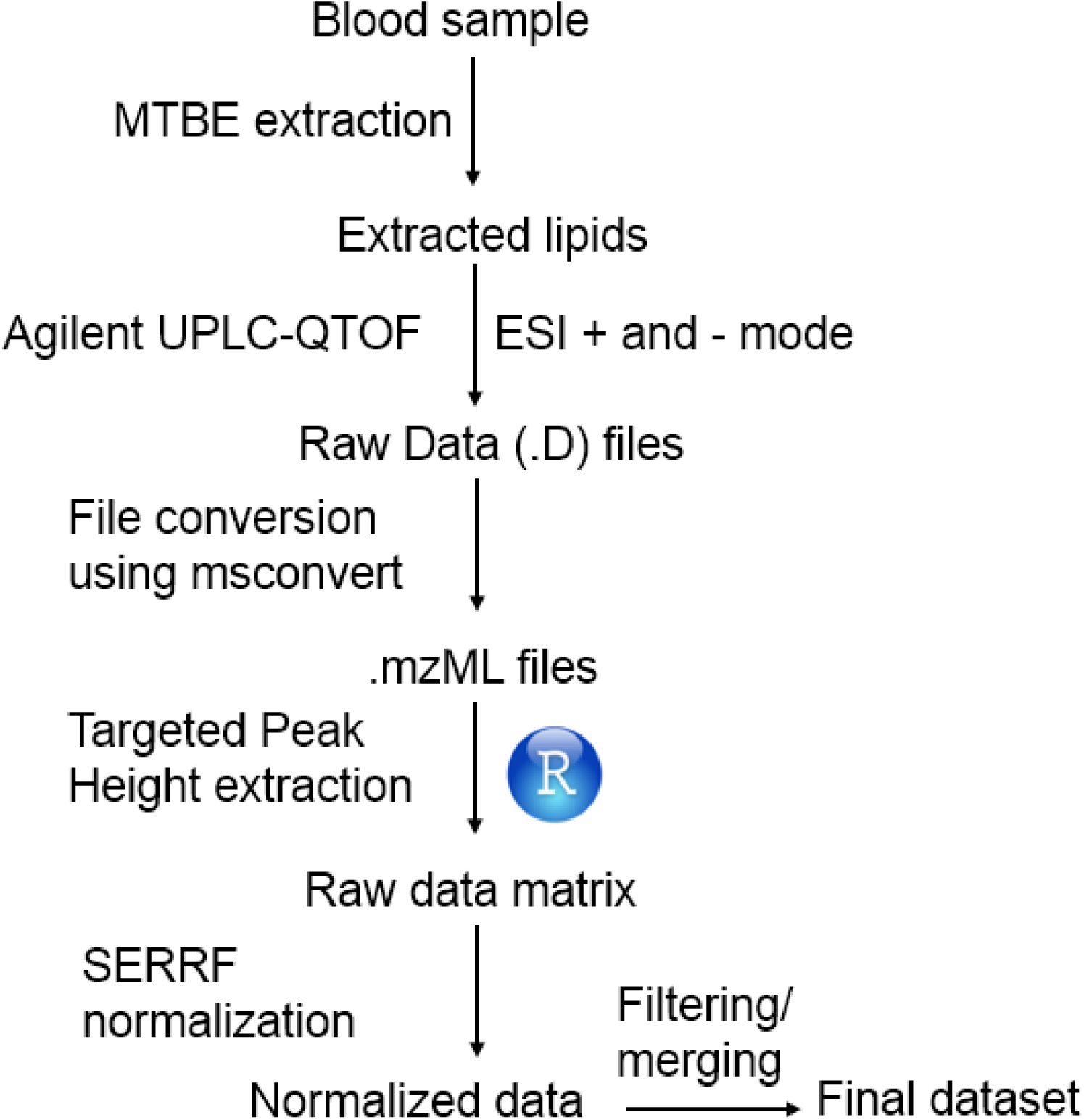
Data processing pipeline overview.

A total 355 unique lipid species covering 15 lipid classes were included in the final dataset, after removing poorly detected and duplicate signals. Random forest-based normalization method using the SERRF tool[12] removed technical variance to as low as 2-6% relative standard deviation across all studies, using BioreclamationIVT plasma QC samples that were analyzed after every 10^th^ sample in all cohorts (see Supplementary Table S1).

### 2.2 Significantly associated individual lipids

Up to 87% of all lipids were found to be significantly different (*p*<0.05) between men and women in at least one cohort study using the raw *p*-values of the Mann-Whitney U test. More lipids were found to be specifically altered in ADNI cohort comparison to the GOLDN and GeneBank cohort studies. 33% of all significantly altered lipids were found to be common across all three studies (Figure 2).

**Figure 2.**
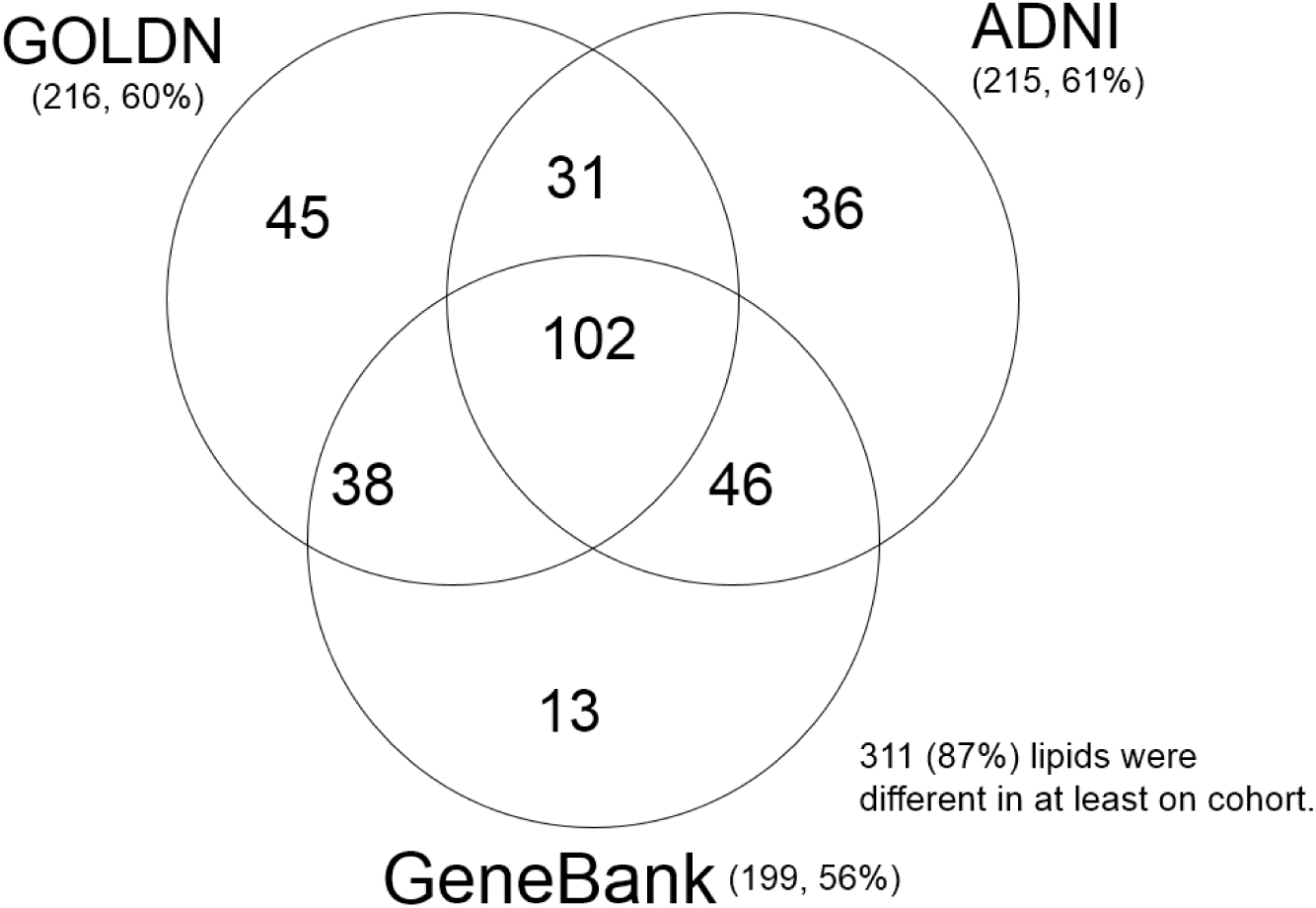
Overlap of significant lipids among three cohorts. 332 (94%) lipids were different in at least on cohort.

Among the top-25 of the most significantly different lipids, several sphingomyelin lipids (SM) were, starting with the most significant lipid SM d32:2 were found at consistently higher levels in women than in men across all cohorts (Table 2-4). Other lipids included monounsaturated free fatty acids (FA), ceramides (Cer), triacylglycerols (TG), lysophosphatidylethanolamines (LPE) and lysophosphatidylcholines (LPC). Statistical results for all lipids are provided in the Table S1.

**Table 2.**
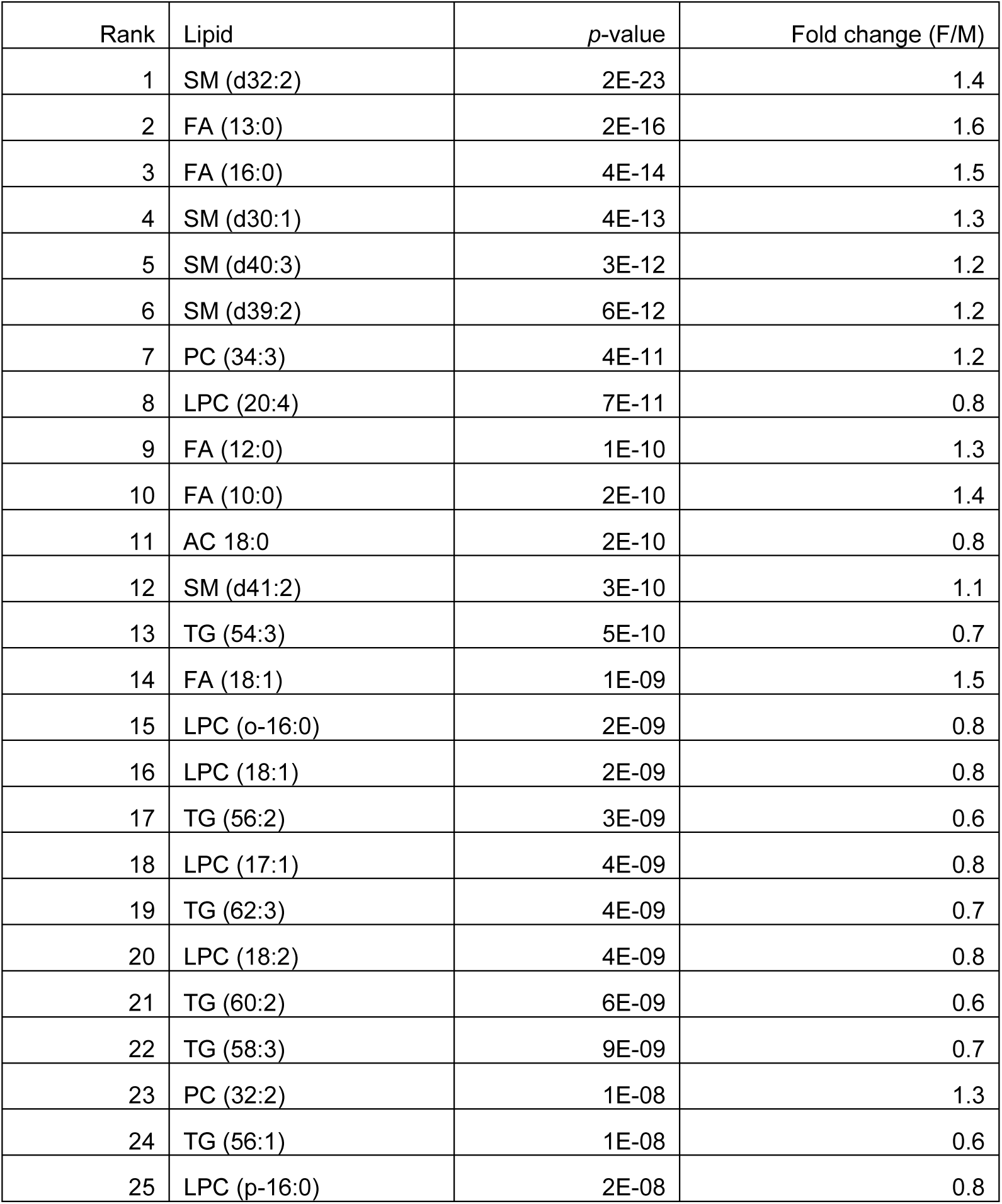
Top 25 significant lipids between male and female in the GOLDN cohort. Fatty acyl groups are annotated by the total number of carbons and the number of double bonds.

| Rank | Lipid | *p*-value | Fold change (F/M) |
| --- | --- | --- | --- |
| 1 | SM (d32:2) | 2E-23 | 1.4 |
| 2 | FA (13:0) | 2E-16 | 1.6 |
| 3 | FA (16:0) | 4E-14 | 1.5 |
| 4 | SM (d30:1) | 4E-13 | 1.3 |
| 5 | SM (d40:3) | 3E-12 | 1.2 |
| 6 | SM (d39:2) | 6E-12 | 1.2 |
| 7 | PC (34:3) | 4E-11 | 1.2 |
| 8 | LPC (20:4) | 7E-11 | 0.8 |
| 9 | FA (12:0) | 1E-10 | 1.3 |
| 10 | FA (10:0) | 2E-10 | 1.4 |
| 11 | AC 18:0 | 2E-10 | 0.8 |
| 12 | SM (d41:2) | 3E-10 | 1.1 |
| 13 | TG (54:3) | 5E-10 | 0.7 |
| 14 | FA (18:1) | 1E-09 | 1.5 |
| 15 | LPC (o-16:0) | 2E-09 | 0.8 |
| 16 | LPC (18:1) | 2E-09 | 0.8 |
| 17 | TG (56:2) | 3E-09 | 0.6 |
| 18 | LPC (17:1) | 4E-09 | 0.8 |
| 19 | TG (62:3) | 4E-09 | 0.7 |
| 20 | LPC (18:2) | 4E-09 | 0.8 |
| 21 | TG (60:2) | 6E-09 | 0.6 |
| 22 | TG (58:3) | 9E-09 | 0.7 |
| 23 | PC (32:2) | 1E-08 | 1.3 |
| 24 | TG (56:1) | 1E-08 | 0.6 |
| 25 | LPC (p-16:0) | 2E-08 | 0.8 |

**Table 3.** Top 25 significant lipids between male and female in the ADNI cohort.

| Rank | Lipid | *p*-value | Fold change (F/M) |
| --- | --- | --- | --- |
| 1 | SM (d32:2) | 5E-52 | 1.40 |
| 2 | SM (d41:2) | 3E-38 | 1.27 |
| 3 | SM (d38:2) | 4E-37 | 1.21 |
| 4 | FA (14:1) | 7E-37 | 1.69 |
| 5 | SM (d39:2) | 7E-33 | 1.38 |
| 6 | FA (16:1) | 8E-33 | 1.81 |
| 7 | PE (38:6) | 2E-31 | 1.51 |
| 8 | SM (d36:2) | 7E-26 | 1.18 |
| 9 | PC (34:3) | 7E-26 | 1.22 |
| 10 | SM (d34:2) | 2E-25 | 1.13 |
| 11 | SM (d30:1) | 8E-25 | 1.30 |
| 12 | HexCer(d34:1(2OH)) | 2E-24 | 1.38 |
| 13 | PC (38:2) | 5E-24 | 1.17 |
| 14 | LPC (18:2) | 8E-24 | 0.80 |
| 15 | SM (d42:3) | 1E-23 | 1.21 |
| 16 | PC (36:6) | 4E-23 | 1.38 |
| 17 | PC (38:7) | 5E-22 | 1.29 |
| 18 | CE (16:1) | 1E-21 | 1.41 |
| 19 | PC (36:5) A | 2E-21 | 1.26 |
| 20 | PC (34:4) | 3E-21 | 1.33 |
| 21 | PC (32:2) | 5E-21 | 1.28 |
| 22 | PE (38:4) | 1E-20 | 1.35 |
| 23 | PC (32:1) | 8E-20 | 1.35 |
| 24 | FA (17:1) | 8E-19 | 1.38 |
| 25 | SM (d40:2) A | 1E-18 | 1.17 |

**Table 4.**
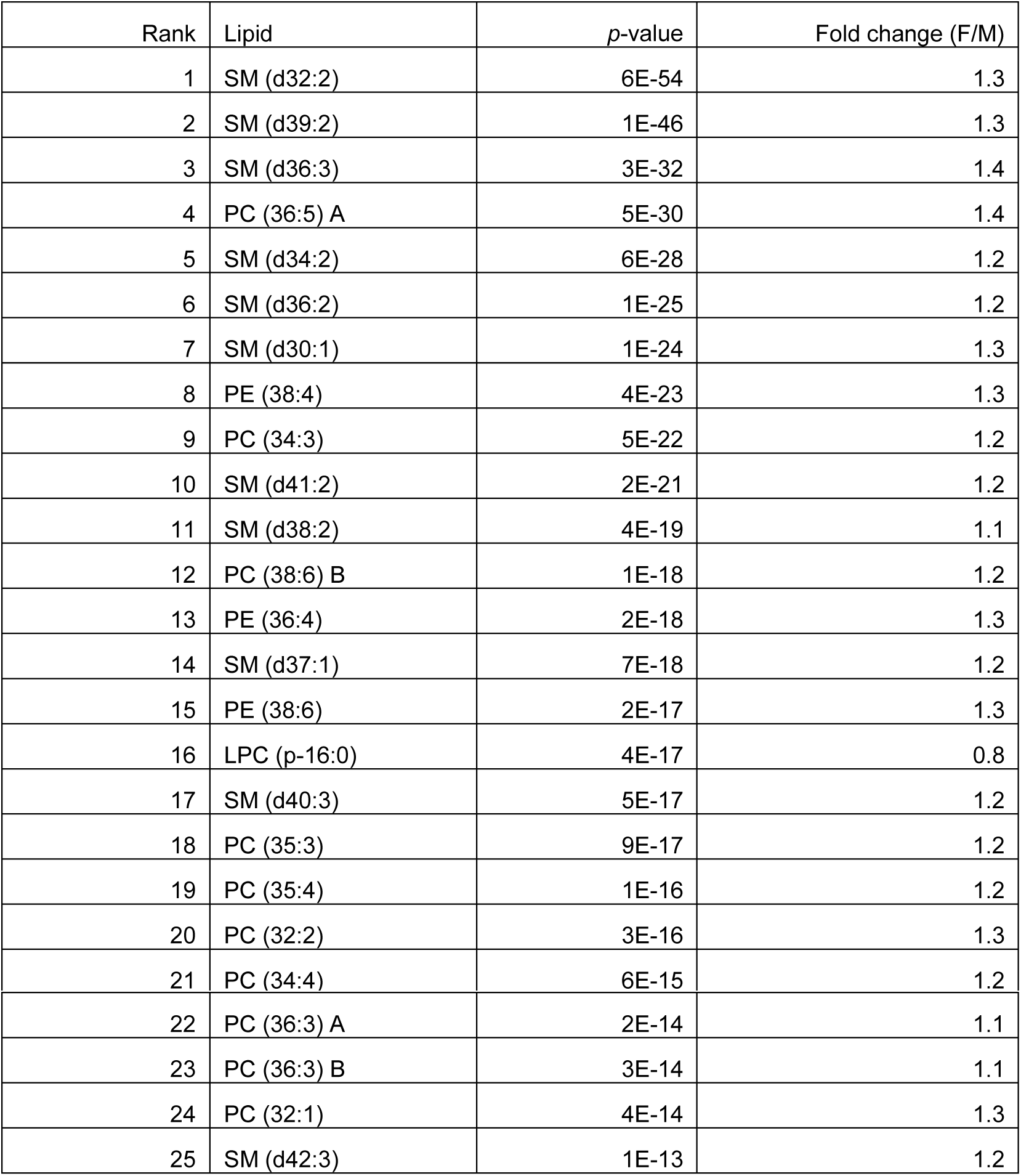
Top 25 significant lipids between male and female in the GeneBank cohort.

### 2.3 Significantly associated lipid classes

Next, we performed a lipid class level analysis to find which chemical classes were significantly higher in the female versus male comparison. We have utilized the ChemRICH enrichment analysis method, which does not rely on a background database for computing the set level statistics. Figure 3 shows the lipids classes associated with differences between both sexes as ChemRICH impact plots.

**Figure 3.**
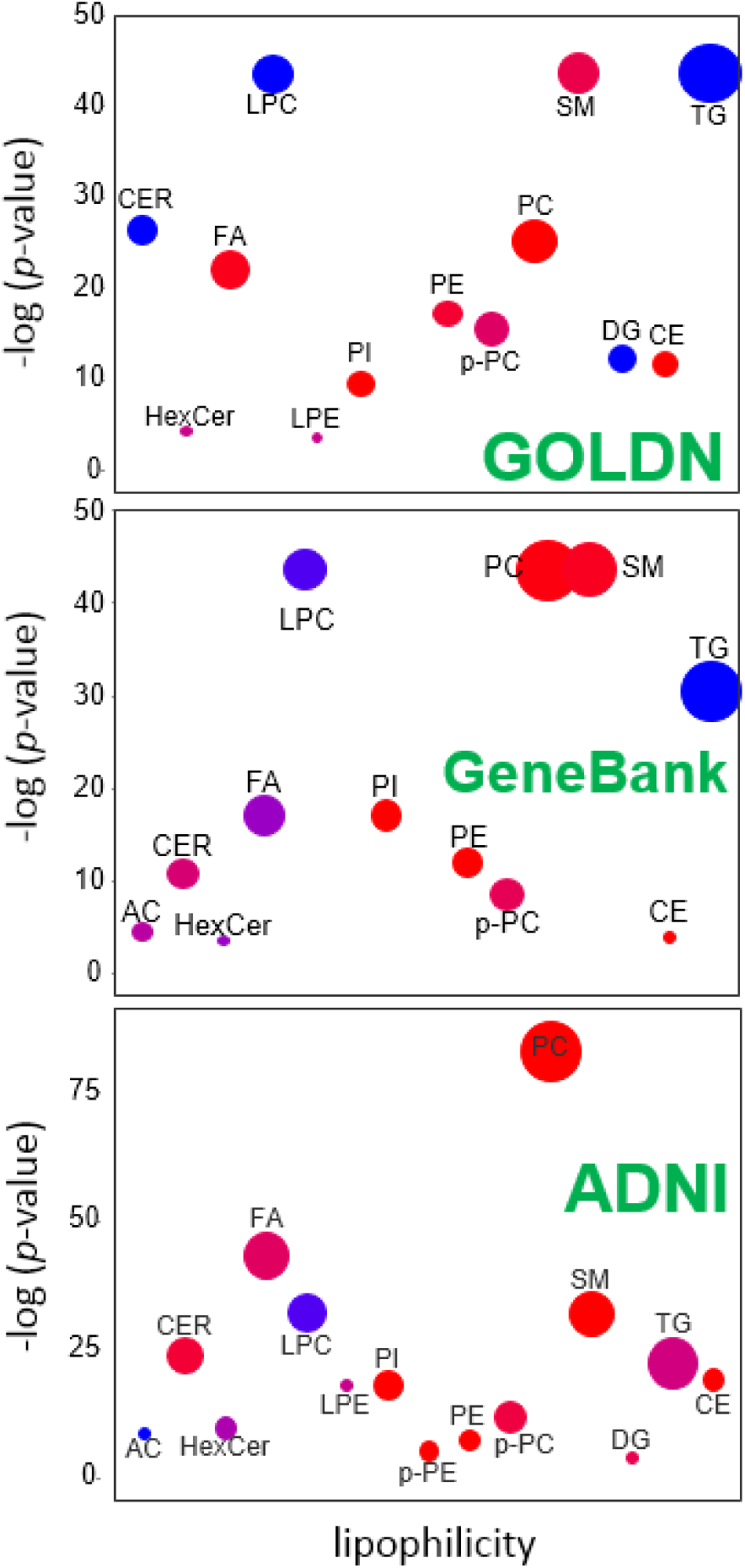
ChemRICH impact plots for the lipid classes associated with sex differences in three cohorts. Red dot means higher in women and blue means higher in men. Purple dot means a mixed response. Size of the dot indicates how many lipids we have in a class. Abbreviations: AC - Acylcarnitine; CE - cholesterol ester; CER - ceramide; DG - diacylglycerol; FA-fatty acid; HexCer - hexosyl ceramide; LPC - Lysophosphatidylcholine; LPE - Lysophosphatidylethanolamine; PC – phosphatidylcholine; PI – phosphatidylcholine; SM - sphinhomyelin; TG - triacylglycerol; p-PC - plasmalogen phosphatidylcholine; p-PE - plasmalogen phosphatidylethanolamine.

All lipid classes were found to be significantly different between men and women in at least one cohort. The most drastic effects were observed for sphingomyelins, triacylglycerol and phosphatidylcholines in the GOLDN and the GeneBank cohorts. Triacylglycerols and lysophosphatidylcholines were consistently higher in men across the three cohorts. Ceramides were higher in men in the GOLDN study but not in the ADNI cohort.

Acylcarnitines were higher in men in the ADNI cohort, but a mixed response was observed for this lipid class in the GeneBank cohort study. Free fatty acids, phosphatidylcholines, sphingomyelins, phosphatidylethanolamine, phosphatidylinositols and cholesterol esters were consistently higher in women in three cohorts. It seems that age was factor for some classes – for example triacylglycerols were lower in women in the younger cohorts (GOLDN and the GeneBank cohort) but not in the ADNI cohort study that included predominately elderly subjects. Interestingly, free fatty acids showed a mixed direction as indicated by purple color in the ChemRICH plot, so we investigated the degree of saturation within this lipid class. We found that saturated fatty acids were higher in men while unsaturated fatty acids were higher in women in all three cohorts.

## 3. Discussion

In comparison to earlier metabolomics studies, we have expanded the sexual dimorphism analysis of blood lipidome with using a comprehensive panel and larger studies. Several new lipids classes were found to be under strong impact of sexual dimorphism. We have replicated the previous finding that levels of sphingomyelins and phosphatidylcholines were elevated in women [9,10] while lysophosphatidylcholines and acylcarnitines were found at higher concentrations in men[6,11]. We also observed differences in triacylglycerol levels in the ADNI study between women and men that could possibly be due to biological age[9], while these differences was absent in comparatively younger participants of the GOLDN and the GeneBank cohort. The most significant lipid clusters included SM, PC, TG, p-PC, LPC and FA lipids in all three cohorts. We found new sex-regulated classes including phosphatidylinositols, plasmalogens, and ceramides. Plasmalogen biosynthesis has been linked with male fertility [13]. It has been previously shown that the hepatic ceramide biosynthesis is regulated by sex hormones, including testosterone [14]. Differences in lipids between both sexes may suggest that sex specific remodeling of lipid metabolism is a fundamental biological process and calls for further studies to discover the underlying mechanisms that can create a basis for developing sex specific disease prevention strategies.

A schema of different metabolic pathways for complex lipids is shown in Figure 4. Our results show a consistently higher ratio of PC to LPC lipids in women compared to men. This ratio likely indicates a higher activity of phospholipases in men to cleave fatty acyl groups from PC membrane lipids to LPC lipid species. We also found higher levels of PC and SM lipids in women, along with lower amounts of ceramides. These three lipid classes intersect in their biochemical pathway and may directly support the idea of higher sphingomyelin synthase activity in women (Figure 4).

Both phospholipases and sphingomyelin synthase are discussed as master regulators of lipid metabolism [15-17]. These enzyme activities could be validated in animal models to identify the specific organs that are contributing to the sexual dimorphism of lipid differences, using functional genomics analysis to construct a sex-specific metabolic network and to map regulatory mechanisms. Combining metabolomics, lipidomics and genomics assays in follow-up studies should be used to further discern the relative contributions from endogenous lipid remodeling versus contributions from diet and exercise.

**Figure 4.**
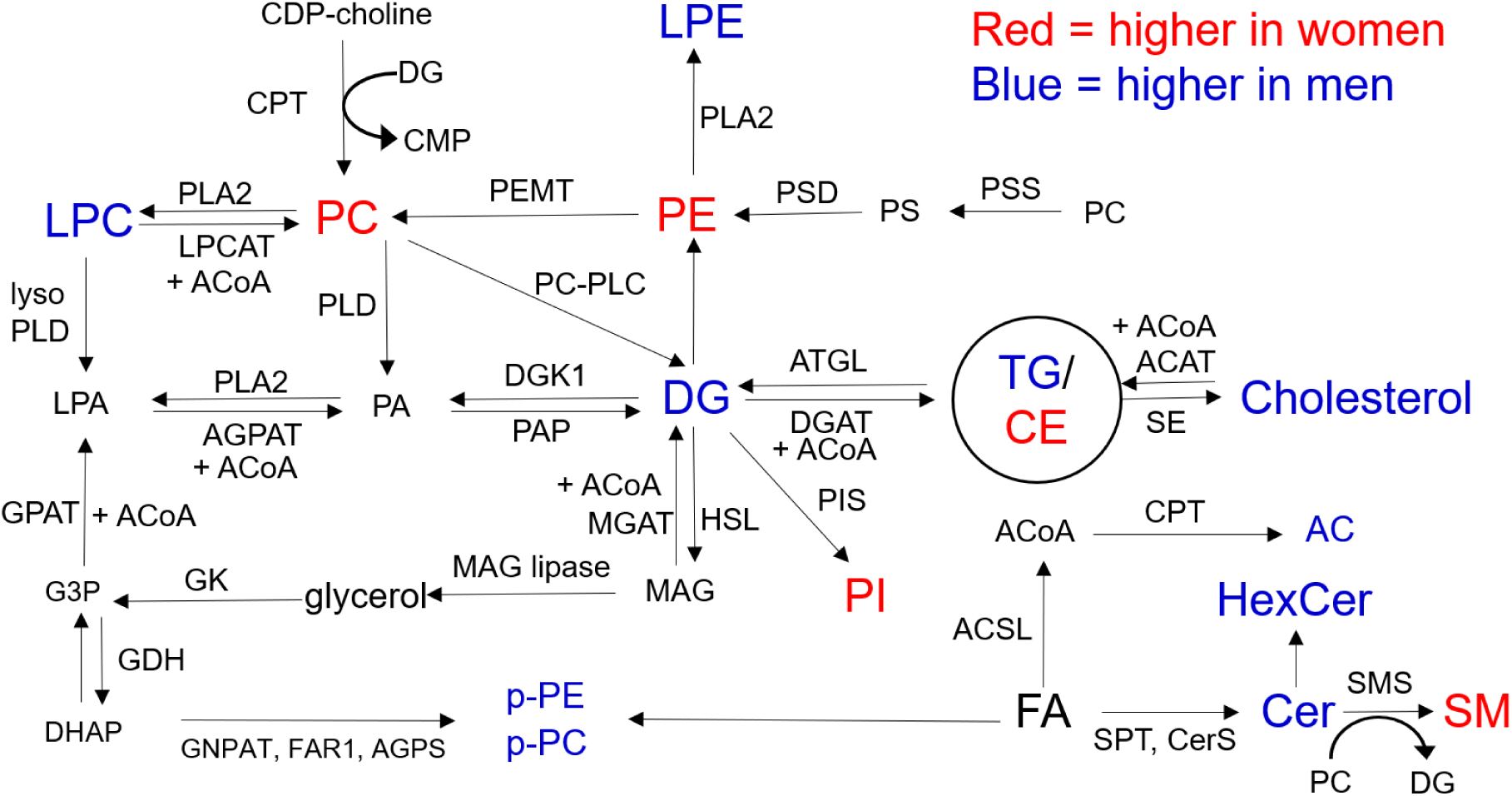
Major metabolic pathways for lipids. Blue indicates higher in male and red indicate higher in female individuals. Abbreviations: AC - Acylcarnitine; ACAT - Acyl-coA:cholesterol o-acyltransferase; ACoA - acyl coenzyme A; AGPS - Alkylglycerone Phosphate Synthase; ATGL - Adipose Triglyceride Lipase; CPT1 - Carnitine Palmitoyltransferase 1A; CDP - cytidine diphosphate; CE - cholesterol ester; CER - ceramide; CMP - cytidine monophosphate; CPT - CDP-choline:1,2-diacylglycerol cholinephosphotransferase; CerS ceramide synthase; DG - diacylglycerol; DGAT - Diacylglycerol *O*-Acyltransferase; DGK1 Diacylglycerol kinase; DHAP - Dihydroxyacetone phosphate; FA-fatty acid; FAR1 - Fatty Acyl-CoA Reductase 1; G3P - Glyceraldehyde 3-phosphate; GDH - Glycerol 3-phosphate dehydrogenase; GK - glycerol kinase; GNPAT - glyceronephosphate O-acyltransferase; GPAT - Glycerol-3-Phosphate Acyltransferase; HSL - Hormone-sensitive Lipase; HexCer hexosyl ceramide; LPA - lysophosphatidic acid; LPC - Lysophosphatidylcholine; LPCAT Lysophosphatidylcholine acyltransferase; LPE - Lysophosphatidylethanolamine; MGAT - Acyl-CoA:monoacylglycerol acyltransferase; PA - phosphatidic acid; PC - phosphatidylcholine; PC-PLC - phospholipase C; PEMT - Phosphatidylethanolamine *N*- Methyltransferase; PI - phosphatidylcholine; PIS - phosphatidylcholine synthase; PLA2 - phospholipase A; PLD - phospholipase D; PS - phosphatidylserine; PSD - Phosphatidylserine decarboxylase; PSS - Phosphatidylserine synthase; SE – sterol esterase; SM - sphinhomyelin; SMS - sphingomyelin synthase; SPT - serine palmitoyl transferase; TG - triacylglycerol; p-PC - plasmalogen phosphatidylcholine; p-PE - plasmalogen phosphatidylethanolamine

The strong sex effects observed in our study suggests that in epidemiological studies, statistical analyses for lipidomics data should considered separately for men and women. We argue against using sex as a confounding or co-variate for regression models to mask the sex-regulated biology. Instead results should be interpreted separately for male and female participants. These drastic sex-regulated differences in the blood lipidome have implications on large epidemiological studies to identify the lipids that can be risk factors for chronic and aging associated disorders. We have observed different patterns for men and women of utilization of saturated and unsaturated free fatty acids, but further studies are needed on remodeling of acyl chains within specific classes of complex lipid[18].

We have looked into three cohorts that have differences in terms of participant’s age, comorbidities, diet and medication usage. Variations in these factors may affect the lipid differences that are associated with sex. One has to consider the impact of monitored, and non-monitored differences in the cohorts examined on the observed results, and additional studies and verification are required. Even though, we have observed remarkable p-values for few sphingomyelins, the sex-effect for these lipids should be checked in other epidemiological cohorts. Nonetheless, our study argue that sex specific differences should be considered in future lipidomic examinations during both study design, and analyses.

## 4. Materials and Methods

### Cohort 1

The GOLDN study (NCT00083369) focuses on how genetic factors interact with environmental (diet and drug) factors to influence triglycerides and other atherogenic lipid species and inflammation markers in blood. The study participants were primarily from three-generational pedigrees from two NHLBI Family Heart Study (FHS) field centers (Minneapolis, MN and Salt Lake City, UT). GOLDN study protocol was approved by the Institutional Review Boards at the University of Minnesota, University of Utah, Tufts University/New England Medical Center, and the University of Alabama at Birmingham[19]. This study included 422 GOLDN participants who were in the extreme tertiles of the lipid response to the high fat diet intervention and had lipidomics data available. The diet intervention has been described [20].

### Cohort 2

The P20 functional metabolomics of cardiovascular disease study consisted of 1,015 participants from the GeneBank cohort, a large (*n* < 10,000) and well-characterized longitudinal tissue repository with associated clinical database at the Cleveland Clinic[21]. All participants gave informed consent and the study was approved by the Cleveland Clinic institutional review board.

### Cohort 3

ADNI is landmark prospective study on Alzheimer’s disease by National Institute of Aging. The study provides imaging, molecular, clinical and neurological function datasets for enrolled subjects along with biospecimens. In this paper, lipidomics data for the baseline serum samples from ADNI-1 study were utilized [22]. The data are available at (http://adni.loni.ucla.edu/). Prior Institutional Review Board approval was obtained at each participating institution and written informed consent was obtained for all participants. Information about the ADNI project is provided on http://www.adni-info.org/.

### Lipid extraction

Lipids were extracted from 20 µL of plasma or serum samples. 225 µL cold methanol containing a mixture of 15 deuterated or odd-chain lipid internal standards was added and samples were vortexed for 10 s. After adding 750 µL of MTBE, samples were vortexed for 10 s and shaken for 5 min at 4°C. Next, 188 µL water was added and samples were vortexed for 20 s and centrifuged for 2 min at 14,000x *g*. One 350 µL aliquots from the non-polar layer was evaporated to dryness in a SpeedVac concentrator. Dried extracts were resuspended using a mixture of methanol/toluene (9:1, v/v) (60 µL) containing an internal standard [12-[[(cyclohexylamino)carbonyl]amino]-dodecanoic acid (CUDA)] used as a quality control. Method blanks and pooled human plasma (BioreclamationIVT) were prepared along with the study samples for monitoring the data quality.

### LC-MS data acquisition

Samples were analyzed using an Agilent 1290 Infinity UHPLC/6530 QTOF MS or an Agilent 1290 Infinity UHPLC/6550 QTOF MS. A charged surface hybrid (CSH) column C18 2.1×100 mm, 1.7 μm column with a VanGuard CSH pre-column, C18 2.1×5 mm, 1.7 μm (both Waters, Milford, MA) were used to separate the extracted lipids. A reference solution of purine and HP-0921 (*m*/*z* 121.0509, *m*/*z* 922.0098 in electrospray ionization (ESI) (+) and *m*/*z* 119.0360 and *m*/*z* 980.0164 (acetate adducts) in ESI(–)) was used to correct small mass drifts during the acquisition. Mobile phase A (60:40 ACN:water + 10 mM ammonium formate + 0.1% formic acid) was prepared by mixing 600 mL ACN, 400 mL water, 1 mL formic acid and 630 mg of ammonium formate. Mobile phase B solvent (90:10 IPA:ACN + 10 mM ammonium formate + 0.1% formic acid) was prepared by mixing 900 mL IPA, 100 mL acetonitrile, 1 mL formic acid, 630 mg ammonium formate previously dissolved in 1 mL of H_2_O. Both solvents were mixed and sonicated for 10 min (twice) before their use. For ESI (–) the composition of mobile phases was identical but 10 mM ammonium acetate (771 mg per 1 L) was used instead as modifier for the ADNI and GOLDN cohort samples. The quadrupole/time-of-flight (QTOF) mass spectrometers were operated with electrospray ionization (ESI) performing full scan in the mass range m/z 100–1700 in positive and negative modes. Instrument parameters were as follows for the ESI (+) mode on the Agilent 6530 QTOF – gas temperature 325 °C, gas flow 8L/min, nebulizer 35 psig, sheath gas temperature 350 °C, sheath gas flow 11, capillary voltage 3500 V, nozzle voltage 1000 V, and fragmentor voltage 120 V. In negative ion mode (Agilent 6550 QTOF), gas temperature 200°C, gas flow 14L/min, fragmentor 175 V, with the other parameters identical to positive ion mode. Data were collected in centroid mode at a rate of 2 scans/s. Injection volume was 1.7 μL for the positive mode and 5 μL for the negative mode. The liquid chromatography gradient used a 0.6 mL/min linear velocity flow rate. The gradient started at 15% B, ramped to 30% at 2 min, 48% at 2.5 min, 82% at 11 min, 99% at 11.5 min and kept at 99% B until 12 min before ramping down to 15% B at 12.1 min which was kept isocratic until 15 min to equilibrate the column. The total run time was 15 min. Samples were analyzed in in multiple batches with batch size ranging from 200-300 samples. After every ten cohort samples, one BioreclamationIVT pooled plasma QC sample was analyzed.

### Targeted signal extraction and data generation

Raw LC-MS data files were converted to the mzML format using the Proteowizard MS Convert utility. These files were imported in R using the mzR package. A database of validated lipids that were routinely detected in blood samples has been compiled at the West Coast Metabolomics Center over the past seven years. In this database, annotated lipids are associated with retention time, adducts, *m*/*z* value and InChI keys, verified by accurate mass, isotope ratio, retention time and MS/MS spectra matching to either commercial lipid standards or to the LipidBlast library[23]. In the database, ESI positive mode had 515 RT-*m*/*z* values and ESI negative mode had 457 RT-*m*/*z* values [22]. Using the retention time–mass-to-charge ratio, RT-*m*/*z* database as a targeted list, we extracted the ion chromatograms (EICs) for all *m*/*z* values using R software with a targeted signal extraction strategy, stretching a range of ±0.5 min of the target retention time for each *m*/*z* value and to obtain the peak height values for all lipids. No peak smoothing or integration method was applied to the EICs. To address retention time drifts between cohorts and among all samples of each cohort study, retention times of the internal standards were obtained for each sample using a ± 0.2 min retention time window to find the peak apexes of the internal standards. A polynomial regression of second order (quadratic) curve was fitted between the expected and observed retention times of the internal standards. The retention times of the remaining compounds in the target database were recalibrated using the regression model. The updated retention times were used with a window of 0.15 min for each metabolite to extract the *m*/*z* intensities values belonging to that ion. Maximum intensity values within the retention time window were used as the peak height of the target compounds. Data were normalized using quality control pool samples that were interjected between every 10 subject samples during data acquisition. We employed the SERRF random forest machine learning algorithm [12] to remove batch and drift effects in each cohort data set for each individual lipid.

### Statistical analysis

Data were log transformed before statistical testing. The Mann-Whitney-U test was used to find raw significance values for each lipid between men and women in each of the three cohorts. *p*-values from the Mann-Whitney-U test were used as input for the chemical similarity enrichment analysis using the ChemRICH software[24] to find statistical significance levels on the basis of lipid classes. ChemRICH *p*-values were corrected for the false discovery rate.

## 5. Conclusions

Our study is the largest lipidomics study to report differences between male and female participants. We have acquired LC-MS datasets on identical mass spectrometers and have used machine learning methods-based signal correction approach to remove technical variance and batch effects. Use of a database independent lipid set enrichment analysis methods have identified a number of specific lipid classes that were associated with differences between adult men and women. Epidemiological studies focusing on these drastically different lipid classes need to stratify the cohort data and interpret the results separately for male and female participants.

## Supplementary Materials

The following are available online at www.mdpi.com/xxx/s1, Table S1: Measured lipids and their statistical significance across three cohorts.

## Author Contributions

D.K.B and O.F. designed the analysis. D.K.B. Y.Z. S.F processed LC/MS data and performed statistical analysis. S.L.H., W.H.W.T., M.R.I., D.K.A and R.K.D. provided blood samples. O.F., T.C. and T.K. acquired lipidomics data. All author contributed in the manuscript writing and approved the content.

## Funding

This work was funded through NIH award U54AI138370, U19AG023122 and U2C ES030158 (OF). The GeneBank study was partially supported by NIH grants DK106000, HL103866, HL076491, HL128300 and HL126827. S.L.H. reports also being supported in part by a Leducq Foundation award, and the Leonard Krieger endowment for Preventive Cardiology. The GOLDN study was supported by NIH R01 HL091357, NIH R01 HL104135, NIH U01 HL072524 and American Heart Association grant 15SDG25760020. The ADNI study was funded through NIH awards U54AI138370 and U19AG023122. National Institute on Aging (R01AG046171, RF1AG051550, and RF1AG057452 and 3U01AG024904-09S4) supported the Alzheimer Disease Metabolomics Consortium which is a part of NIA national initiatives AMP-AD and M2OVE AD. Data collection and sharing for this project was funded by the Alzheimer’s Disease Neuroimaging Initiative (ADNI) (National Institutes of Health Grant U01 AG024904) and DOD ADNI (Department of Defense award number W81XWH-12-2-0012).

## Conflicts of Interest

Dr. Hazen reports being named as co-inventor on pending and issued patents held by the Cleveland Clinic relating to cardiovascular diagnostics or therapeutics, and having the right to receive royalty payment for inventions or discoveries related to cardiovascular diagnostics or therapeutics. Dr. Hazen also reports having been paid as a consultant for P&G, and receiving research funds from Astra Zeneca, P&G, Pfizer Inc., and Roche Diagnostics.

